# Still poor HAART adherence has great impact on HIV/AIDS treatment failure in Ethiopia

**DOI:** 10.1101/440743

**Authors:** Aklilu Endalamaw, Mengistu Mekonen, Demeke Debebe, Fekadu Ambaw, Hiwot Tesera, Tesfa Dejenie Habtewold

## Abstract

**Background:** The pooled burden of HIV treatment failure and its associated factors in Ethiopian context is required to provide evidence towards renewed ambitious future goal.

**Methods:** Ethiopian Universities’ (University of Gondar and Addis Ababa University) online repository library, Google scholar, PubMed, Web of Science, and Scopus were used to get the research articles. I-squared statistics was used to see heterogeneity. Publication bias was checked by Egger’s regression test. The DerSimonian-Laird random effects model was employed to estimate the overall prevalence. Subgroup analysis based on geographical location of the study, study population by age, treatment failure type, and study design was conducted to see variation in outcomes. The sensitivity analysis was also employed to see whether the outlier result found in the included studies.

**Results:** Overall HIV treatment failure found to be 15.9% (95% CI: 11.6%-20.1%). Using immunological definition, HIV treatment failure was 10.2% (6.9%-13.6%); using virological definition of treatment failure (5.6% (95% CI: 2.9%-8.3%) and clinical definition of treatment failure (6.3% (4.6%-8.0%)) were also determined. The pooled effects of WHO clinical stage III/IV (AOR=1.9; 95% CI: 1.3-2.6), presence of opportunistic infections (AOR=1.8; 95% CI: 1.2-2.4), and poor HAART adherence (AOR= 8.1; 95% CI: 4.3-11.8) on HIV treatment failure are estimated.

**Conclusions:** HIV treatment failure in Ethiopia found to be high. HIV intervention programs need to address the specified contributing factors of HIV treatment failure. Behavioral intervention to prevent treatment interruption is required to sustain HIV treatment adherence.

**Protocol Registration:** It has been registered in the PROSPERO database (CRD42018100254).

## BACKGROUND

The risk of death due to HIV has been decreased after the era of highly active antiretroviral therapy (HAART) (1). Evidence has shown that individuals on HAART with an undetectable viral load, absence of advanced clinical finding, and high CD4 count are less likely to transmit HIV to others people (2, 3). However, the risk of HIV transmission is high due to treatment failure. Treatment failure can be virological, immunological, or clinical failure. Virological failure is plasma viral load above 1000 copies/ ml based on two consecutive viral load measurements after 3 months, with adherence support. Immunological failure is CD4 count falls to the baseline (or below) or persistent CD4 levels below 100 cells/mm3 for adult and adolescent or below 200 cells/mm3 younger than 5 years. Clinical failure is defined as occurrence or recurrence of advanced WHO clinical stage or conditions after at least 6 months of therapy (4). Globally, UNAIDS’ 90-90-90 planned to have 90% of people on HAART are virally suppressed by 2030 and as a result HIV treatment failure would be prevented (5). Despite this ambitious goal, as of a systematic analysis of national HIV treatment cascades of 69 countries by 2016, viral suppression was between 68% in Switzerland and 7% in China (6). It can be prevented through the implementation of globally recommended strategies. For instance, increase HAART adherence, taking medication based on the appropriate prescription, prevent drug-drug interaction, increasing knowledge and attitudes of patients towards HAART, timely initiation of HAART, prevention and control of opportunistic infections, and implementation of effective food and nutrition policy (7-11).

A higher viral load may leads to HIV treatment failure, which is becoming a threat of different African countries, like in Burkina Faso (6.4%) (12), Ghana (15.7%) (13), and Tanzania (14.9%) (14). In Ethiopia, virological, immunological, and clinical failure found to be in the range between 1.3% (15) to 11.5% (16), 2.1% (17) to 21% (18), and 3.1% (19) to 12.3% (20) respectively.

With these variations of reports, there is no pooled representative national data in Ethiopia. In order to provide evidence towards renewed ambitious future goal, it is now critical to reflect the pooled burden of HIV treatment failure in Ethiopian context. The objective of this study was firstly, to estimate the national burden of HIV treatment failure and secondly, to review contextual factors of HIV treatment failure using globally accepted key performance indicators as a framework. Thus, this information will be helpful for healthcare professionals and further helps to enable the country to sustain its successes and identify and improve weaknesses towards the goal of ending AIDS.

## METHODS

### Reporting

It is reported based on the Preferred Reporting Items for Systematic Reviews and Meta-analyses (PRISMA) guideline (21) (supplementary file-research checklist). Its protocol is registered in the Prospero database with a registration number CRD42018100254.

### Search strategy

PubMed, Web of Science, Scopus, and Google Scholar databases were used to get the research articles. The search strategy made in PubMed was: [(“Human Immunodeficiency virus”[MeSH Terms] OR HIV OR AIDS OR “Acquired Immunodeficiency syndrome” AND (“antiretroviral therapy”[MeSH Terms] OR “highly antiretroviral therapy” OR HAART OR ART OR “ARV Therapy” OR “antiretroviral therapy”) AND (outcome OR “treatment failure” OR failure OR “virological failure” OR “immunological failure” OR “Clinical failure”) AND (Ethiopia)]. The search done in PubMed through search terms was 03/10/2018. In addition, Ethiopian Universities’ (University of Gondar and Addis Ababa University) online repository library were searched. Endnote 7 reference manager software was used to manage duplicated references and for citation in the text.

### Inclusion and exclusion criteria

Those articles included in this meta-analysis were: (1) cohort, case-control, and cross-sectional studies, (2) studies that reported the prevalence and/ or AOR (adjusted odd ratio) of associated factors of overall HAART treatment, immunological, clinical, and virological failure, (3) studies conducted in Ethiopia, and (4) studies published in English.

Studies without full-text access, qualitative studies, and conference proceeding without full-text report were excluded.

### Outcome measurement

According to WHO (4), HIV treatment failure could be clinical, immunological, and virological failure.

The prevalence of failure was ascertained by dividing the participants with the outcome of interests to the overall study participants multiplied by 100.

### Quality assessment

Two authors assessed the quality of the articles based on Newcastle-Ottawa Scale quality assessment tool for cross-sectional, case control, and cohort studies (22). The criteria for cross-sectional studies has mainly three sections, in which the first section mainly focused on selection and graded by four stars, the second section dedicated with the comparability of the study and graded by two stars, and the third section emphasized on the outcome and graded by three stars. The criteria for case control studies were: 1) selection evaluated by four stars, 2) comparability assessed by two stars, and 3) exposure graded by four stars. The criteria for cohort studies were: 1) selection graded by six stars, 2) comparability graded by two stars, and 3) outcome graded by five stars. Whenever disagreement happened between the two quality assessors, the procedure would be repeated and further solved by interference of third reviewer. Cross-sectional, case-control, and cohort studies scored 6 and/or above, 7 and/or above, and 9 and/or above quality assessment criteria were included respectively.

### Data extraction process

Two authors extracted the required data. The first author and year of publication, sample size, an outcome of interest, study design, study population, geographical location of the study, fund, and response rate were collected.

### Data synthesis and statistical analysis

STATA 14 (Stata Corp, College Station, TX, USA) statistical software was used for meta-analysis. Publication bias assessed by funnel plot and more objectively by Egger’s regression test. I-squared statistics was used to check the heterogeneity of the studies. The DerSimonian-Laird random-effects model was employed to estimate the overall prevalence. Subgroup analysis based on geographical location of the study, treatment failure type, study population by age, and study design was conducted to see variation in outcomes. The sensitivity analysis was also employed to see whether the outlier result found in the included studies.

## RESULTS

### Search results

A total of 873 articles were found from search engines (Figure 1).

**Figure 1:**
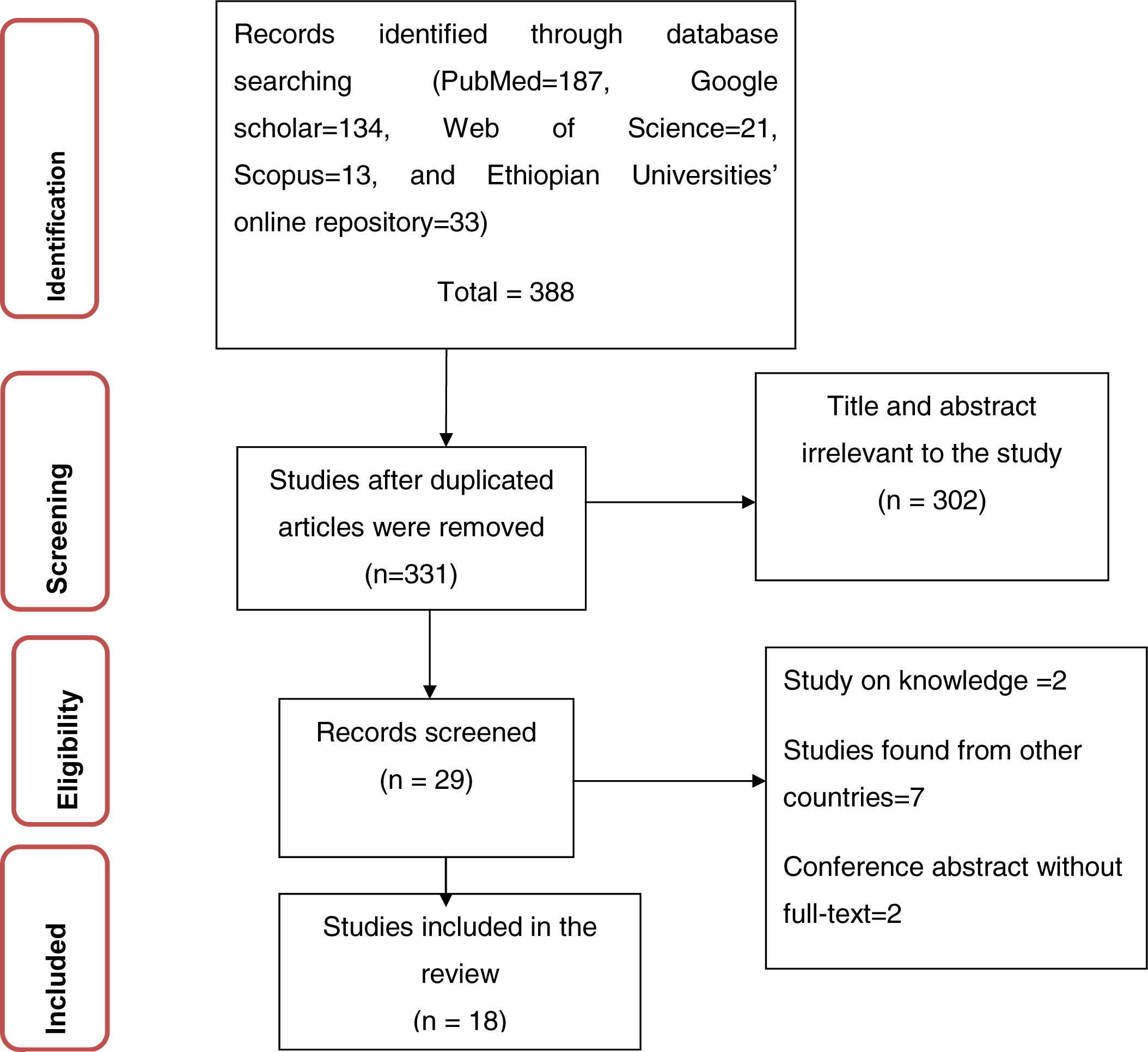
PRISMA flow chart diagram describing selection of studies

### Publication bias

Funnel plot for HIV treatment failure is shown below (Figure 2). Egger’s regression test of p-value for overall HIV treatment failure is 0.226.

**Figure 2:**
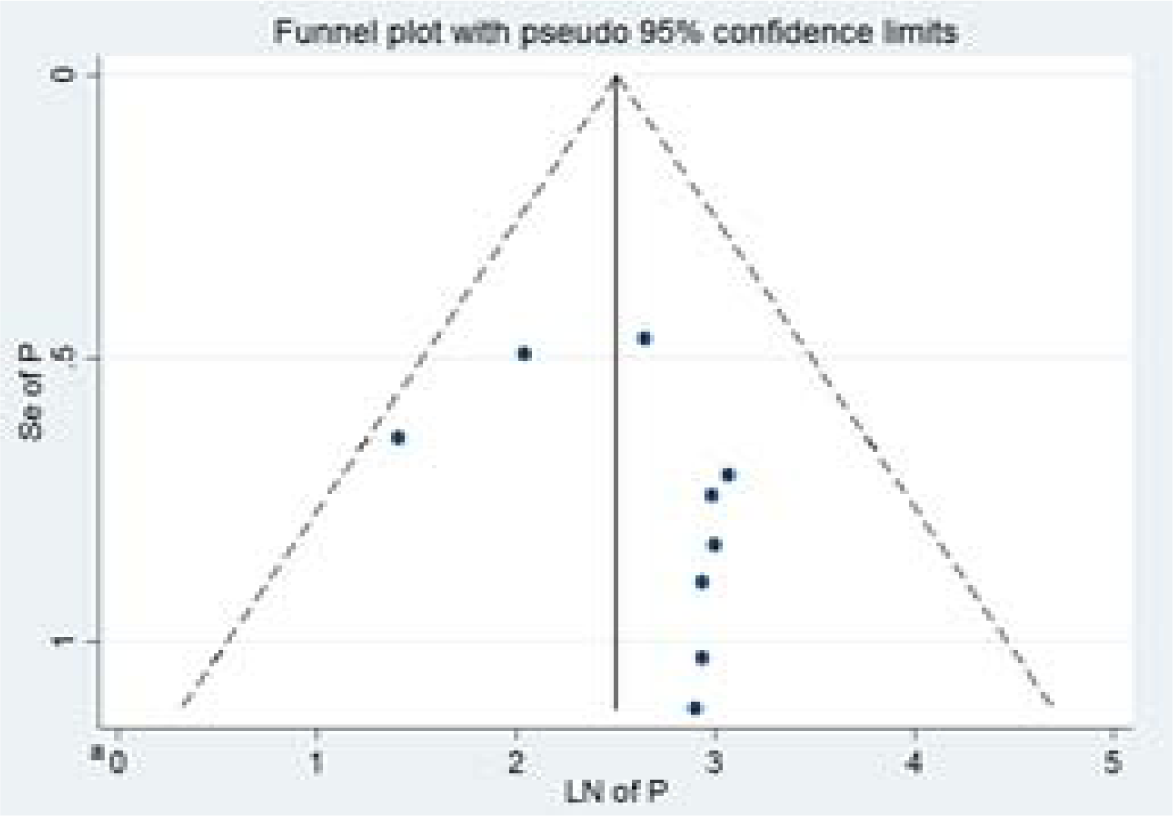
Funnel plot, in which vertical line indicates the effect size whereas diagonal line indicates precision of individual studies with 95% confidence limit

### Meta-analysis

#### HIV treatment failure based on definition of HAART failure

A total of 4,738 participants in nine studies were used to estimate the pooled prevalence of HIV treatment failure based on the definition of HAART failure. The pooled prevalence of HIV treatment failure was 15.9% (95% CI: 11.6%-20.1%) (Figure 3).

**Figure 3:**
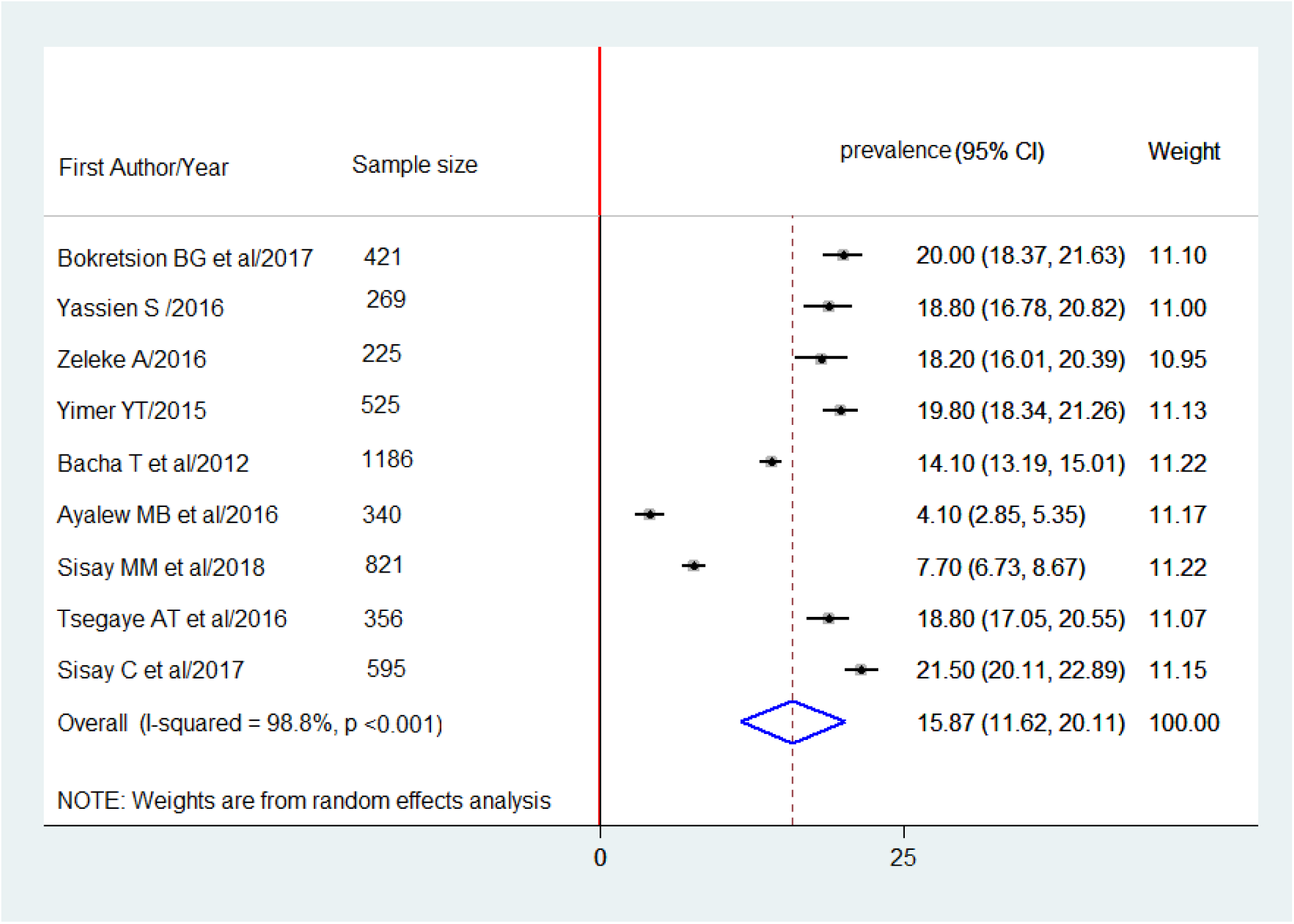
Forest plot of the prevalence of HAART failure in Ethiopia and its 95%CI, the midpoint of each line illustrates the prevalence rate estimated in each study. The diamond shows pooled prevalence.

#### Immunological and Virological definition of HIV treatment failure

A total of 5,899 study participants in 13 studies were involved to determine HIV treatment failure based on immunological definition. Of which, 10.2% (6.9%-13.6%) developed immunological failure. Regarding virological failure, the pooled prevalence from six studies with a total of 2,406 participants was 5.6% (95% CI: 2.9%-8.3%).

#### Clinical definition of HIV treatment failure

A total of 4,497 study participants in 9 studies found to estimate the clinical failure, in which the pooled prevalence was 6.3% (4.6%-8.0%).

#### Subgroup analysis

Subgroup analysis was employed based on region, age of the study participants, and study design. Lower prevalence of HIV treatment failure based on the definition of HAART, immunological, and virological failure is found in Amhara (13.7%), Tigray (6.5%), and Addis Ababa (1.5%) respectively.

#### Sensitivity Analysis

In the sensitivity analysis, the overall HIV treatment failure based on the definition of HAART failure was observed high (17.3%) and low (15.2%) when *Ayalew MB et al 2016* and *Sisay C et al/2017* was omitted respectively.

#### Associated factors of HIV treatment failure

HIV treatment failure is attributed to clinical, drug and health system-related factors.

##### Clinical-related factors

The pooled effects of CD4 cell count <200 cells/mm3 (AOR=7.2; 95% CI: 2.5-12.0), ≤ 100 cells/ mm3 (AOR=2.1; 95% CI: 1.4-2.8) and <50 cells/mm3 (AOR=3.3; 95% CI: 1.4-5.3) as compared to those with >200, >100, and > 50 cells/mm3 on HIV treatment failure were estimated respectively.

The pooled effect of being on WHO clinical stage III/IV showed higher risk (AOR=1.9; 95% CI: 1.3-2.6) to HIV treatment failure as compared to stage II/I. The pooled effect presence of opportunistic infections (TB, diarrhea, pneumonia, other OIs) is more likely (AOR=1.8; 95% CI: 1.2-2.4) to exposed patients to HIV treatment failure.

#### Drug-related factors

The pooled effect of poor HAART adherence on HIV treatment failure was found to be 8.1 (95% CI: 4.3-11.8)

## DISCUSSION

Our study has two main findings related to the national prevalence and risk factors for HIV treatment failure. First, we noted that using the definition of HAART failure, HIV treatment failure was 15.9% (95% CI: 11.6%-20.1%).

In Ethiopia threat of HIV treatment failure is becoming appointed discussion. This might be due to implementation of poor HIV care, delaying on recognition of symptom of treatment failure, (35), late initiation of HAART (36), high burden of opportunistic infections (37), lack of well nutritional support (38), ART associated adverse reaction,(39) and frequent psychological problem (40, 41) are recorded in Ethiopia. Besides, absence of frequent therapeutic drug monitors and/ or resistance testing while the patient is still on the suspect or failing regimen.

Though it is found that the WHO immunological criteria have a very low sensitivity and high specificity (42), this finding showed that HIV treatment failure using immunological definition of treatment failure (10.2%) was higher than that of using clinical (6.3%) and virological (5.6%) definition of treatment failure. These variations might be due to the number of studies included to immunological definition of HIV treatment failure were many in number. The lower prevalence of HIV treatment failure using clinical definition might be due to limited diagnostic capabilities. Although only five studies were found to estimate virological failure that might result under-estimation, the third 90 target of UNAIDS seems to be achieved; there is a plan to achieve 90% of all people receiving ART will have viral suppression by 2020 (5).

Based on the subgroup analysis, HIV treatment failure is lower in children. ART monitoring using clinical and immunological criteria is problematic in children, and misclassification rates using the WHO pediatric guidelines remain high (43).

It is estimated that lower CD4 cell count and advance WHO clinical stage leads to HIV treatment failure. Another studies (44, 45) reported similar finding in other settings. The presence of opportunistic infections on the other had linked to CD4 cell level. As patients’ immune status becomes compromised, the rate of viral replication increases. CD4 cell count is the back bone of immunity construction that helps the human body to protect from the disease and can be prevent HIV replication (46).

Poor HAART adherence found to have a great impact on the occurrence of HIV treatment failure. It is widely agreed that once treatment is initiated, it should not be interrupted. In Ethiopia, within 7 days, nearly 11.3% of children are poorly adhered to ART (47). It is expected that as duration increased the probability of ART interruptions would be more likely. The same in adult HIV patients’ treatment interruption is fall in the range between 11.8-25.8% (48, 49). HIV treatment failure in Ethiopia found to be high. The current finding will have health policy and clinical implication for therapeutic management decisions. Early identification of ART treatment failure allows patients a higher chance of success when switching to a second line ART. Report on HIV treatment failure will be used to monitor the progress of the national action plan of 90-90-90 strategies.

## LIMITATION OF THE STUDY

Lack of studies in some geographical areas of Ethiopia and therefore the finding should be interpreted with caution.

## REFERENCES

1. Quinn TC. HIV epidemiology and the effects of antiviral therapy on long-term consequences. AIDS (London, England). 2008;22(Suppl 3):S7.

2. Hull MW, Montaner JS. HIV treatment as prevention: the key to an AIDS-free generation. Journal of food and drug analysis. 2013;21(4):S95–S101.

3. Granich R, Crowley S, Vitoria M, Smyth C, Kahn JG, Bennett R, et al. Highly active antiretroviral treatment as prevention of HIV transmission: review of scientific evidence and update. Current Opinion in HIV and AIDS. 2010;5(4):298.

4. World Health Organization. Consolidated guidelines on the use of antiretroviral drugs for treating and preventing HIV infection. Geneva, Switzerland: 2013.

5. Sidibé M, Loures L, Samb B. The UNAIDS 90–90–90 target: a clear choice for ending AIDS and for sustainable health and development. Journal of the International AIDS Society. 2016;19(1):21133.

6. Levi J, Raymond A, Pozniak A, Vernazza P, Kohler P, Hill A. Can the UNAIDS 90-90-90 target be achieved? A systematic analysis of national HIV treatment cascades. BMJ Global Health. 2016;1(2):e000010.

7. Nachega JB, Adetokunboh O, Uthman OA, Knowlton AW, Altice FL, Schechter M, et al. Community-based interventions to improve and sustain antiretroviral therapy adherence, retention in HIV care and clinical outcomes in low-and middle-income countries for achieving the UNAIDS 90-90-90 targets. Current HIV/AIDS Reports. 2016;13(5):241–55.

8. Organization WH. ART failure and strategies for switching ART regimens in the WHO European Region: report of the WHO expert consultation, Copenhagen, 7 December 2007. 2008.

9. Beisel WR. Nutrition and immune function: overview. The Journal of nutrition. 1996;126(suppl_10):2611S–5S.

10. Haddad L, Gillespie S. Effective food and nutrition policy responses to HIV/AIDS: what we know and what we need to know. Journal of International Development: The Journal of the Development Studies Association. 2001;13(4):487–511.

11. Agu KA, Oparah AC, Ochei UM. Knowledge and attitudes of HIV-infected patients on antiretroviral therapy regarding adverse drug reactions (ADRs) in selected hospitals in Nigeria. Perspectives in clinical research. 2012;3(3):95.

12. Zoungrana J et al. Prevalence and Factors Associated with Treatment Failure during Antiretroviral Therapy Atbobo-Dioulasso University Teaching Hospital (Burkina Faso) (2008-2013). Austin J HIV/AIDS Res 2016;3(2).

13. Owusu M, Mensah E, Enimil A, Mutocheluh M. PREVALENCE AND RISK FACTORS OF VIROLOGICAL FAILURE AMONG CHILDREN ON ANTIRETROVIRAL THERAPY. BMJ Global Health. 2017;2(Suppl 2):A9–A.

14. Hawkins C, Ulenga N, Liu E, Aboud S, Mugusi F, Chalamilla G, et al. HIV virological failure and drug resistance in a cohort of Tanzanian HIV-infected adults. Journal of Antimicrobial Chemotherapy. 2016;71(7):1966–74.

15. Yimer YT, Yalew AW. Magnitude and predictors of anti-retroviral treatment (ART) failure in private health facilities in Addis Ababa, Ethiopia. PLoS One. 2015;10(5):e0126026.

16. Hailu GG, Hagos DG, Hagos AK, Wasihun AG, Dejene TA. Virological and immunological failure of HAART and associated risk factors among adults and adolescents in the Tigray region of Northern Ethiopia. PLoS one. 2018;13(5):e0196259.

17. Sisay MM, Ayele TA, Gelaw YA, Tsegaye AT, Gelaye KA, Melak MF. Incidence and risk factors of first-line antiretroviral treatment failure among human immunodeficiency virus-infected children in Amhara regional state, Ethiopia: a retrospective follow-up study. BMJ open. 2018;8(4):e019181.

18. Yayehirad AM, Mamo WT, Gizachew AT, Tadesse AA. Rate of immunological failure and its predictors among patients on highly active antiretroviral therapy at Debremarkos hospital, Northwest Ethiopia: a retrospective follow up study. Journal of AIDS and Clinical Research. 2013;4(5).

19. Tsegaye AT, Wubshet M, Awoke T, Alene KA. Predictors of treatment failure on second-line antiretroviral therapy among adults in northwest Ethiopia: a multicentre retrospective follow-up study. BMJ open. 2016;6(12):e012537.

20. Yassin S, Gebretekle GB. Magnitude and predictors of antiretroviral treatment failure among HIV-infected children in Fiche and Kuyu hospitals, Oromia region, Ethiopia: a retrospective cohort study. Pharmacology research & perspectives. 2017;5(1):e00296.

21. Liberati A, Altman DG, Tetzlaff J, Mulrow C, Gøtzsche PC, Ioannidis JP, et al. The PRISMA statement for reporting systematic reviews and meta-analyses of studies that evaluate health care interventions: explanation and elaboration. PLoS medicine. 2009;6(7):e1000100.

22. Wells GA, Shea, B., O’Connell, D., Peterson, J., Welch, V., et al,. The Newcastle-Ottawa Scale (NOS) for assessing the quality of nonrandomized studies in meta-analysis 2011.

23. Bokretsion GB, Endalkachew N, Getachew KA. HIV/AIDS treatment failure and its determinant factors among first line HAART patients at Felege-Hiwot Referral Hospital, Bahir Dar, Northwest Ethiopia. Journal of AIDS and Clinical Research. 2017;8(11).

24. Zeleke A. Prevalence of antiretroviral treatment failure and associated factors in HIV infected children on antiretroviral therapy at Gondar University Hospital, retrospective cohort study. International Journal of Medicine and Medical Sciences. 2016;8(11):125–32.

25. Ayalew MB, Kumilachew D, Belay A, Getu S, Teju D, Endale D, et al. First-line antiretroviral treatment failure and associated factors in HIV patients at the University of Gondar Teaching Hospital, Gondar, Northwest Ethiopia. HIV/AIDS (Auckland, NZ). 2016;8:141.

26. Babo YD, Alemie GA, Fentaye FW. Predictors of first-line antiretroviral therapy failure amongst HIV-infected adult clients at Woldia Hospital, Northeast Ethiopia. PLoS one. 2017;12(11):e0187694.

27. Bayu B, Tariku A, Bulti AB, Habitu YA, Derso T, Teshome DF. Determinants of virological failure among patients on highly active antiretroviral therapy in University of Gondar Referral Hospital, Northwest Ethiopia: a case-control study. HIV/AIDS (Auckland, NZ). 2017;9:153–9.

28. Teshome W, Assefa A. Predictors of immunological failure of antiretroviral therapy among HIV infected patients in Ethiopia: a matched case-control study. PLoS One. 2014;9(12):e115125.

29. Bacha T, Tilahun B, Worku A. Predictors of treatment failure and time to detection and switching in HIV-infected Ethiopian children receiving first line anti-retroviral therapy. BMC infectious diseases. 2012;12(1):197.

30. Sisay C, Bekele A, Sisay A, Mekonen H, Terfa K. Incidence and Predictors of Anti-Retroviral Treatment (ART) Failure among Adults Receiving HIV Care at Zewditu Memorial Hospital, Addis Ababa, Ethiopia. J AIDS Clin Res. 2017;8(749):2.

31. Getnet Y. Determinants of First Line Antiretroviral Treatment Failure in Public Hospitals of Addis Ababa, Ethiopia: Unmatched Case Control Study. Journal of Biology, Agriculture and Healthcare. 2014;4(224-3208):1–12.

32. Abdissa A, Yilma D, Fonager J, Audelin AM, Christensen LH, Olsen MF, et al. Drug resistance in HIV patients with virological failure or slow virological response to antiretroviral therapy in Ethiopia. BMC infectious diseases. 2014;14(1):181.

33. Workneh N, Girma T, Woldie M. Immunologic and clinical outcomes of children on HAART: a Retrospective cohort analysis at Jimma University specialized hospital. Ethiopian Journal of Health Sciences. 2009;19(2).

34. Tadesse BT, Foster BA, Jerene D, Ruff A. Cohort profile: improving treatment of HIV-infected Ethiopian children through better detection of treatment failure in southern Ethiopia. BMJ open. 2017;7(2):e013528.

35. Deribew A, Biadgilign S, Berhanu D, Defar A, Deribe K, Tekle E, et al. Capacity of health facilities for diagnosis and treatment of HIV/AIDS in Ethiopia. BMC health services research. 2018;18(1):535.

36. Nash D, Tymejczyk O, Gadisa T, Kulkarni SG, Hoffman S, Yigzaw M, et al. Factors associated with initiation of antiretroviral therapy in the advanced stages of HIV infection in six Ethiopian HIV clinics, 2012 to 2013. Journal of the International AIDS Society. 2016;19(1):20637.

37. Haileamlak A, Hagos T, Abebe W, Abraham L, Asefa H, Teklu AM. Predictors of Hospitalization among Children on ART in Ethiopia: a Cohort study. Ethiopian journal of health sciences. 2017;27(1):53–62.

38. Gebremichael DY, Hadush KT, Kebede EM, Zegeye RT. Food Insecurity, Nutritional Status, and Factors Associated with Malnutrition among People Living with HIV/AIDS Attending Antiretroviral Therapy at Public Health Facilities in West Shewa Zone, Central Ethiopia. BioMed research international. 2018;2018.

39. Abdissa S, Fekade D, Feleke Y, Seboxa T, Diro E. Adverse drug reactions associated with antiretroviral treatment among adult Ethiopian patients in a tertiary hospital. Ethiopian medical journal. 2012;50(2):107–13.

40. Selamawit Z, Nurilign A. Common mental disorder among HIV infected individuals at Comprehensive HIV Care and Treatment Clinic of Debre Markos referral Hospital, Ethiopia. Journal of AIDS and Clinical Research. 2015;6(2).

41. Amare T, Getinet W, Shumet S, Asrat B. Prevalence and Associated Factors of Depression among PLHIV in Ethiopia: Systematic Review and Meta-Analysis, 2017. AIDS research and treatment. 2018;2018.

42. Le NK, Riggi E, Marrone G, Van Vu T, Izurieta RO, Nguyen CKT, et al. Assessment of WHO criteria for identifying ART treatment failure in Vietnam from 2007 to 2011. PLoS one. 2017;12(9):e0182688.

43. Westley BP, DeLong AK, Tray CS, Sophearin D, Dufort EM, Nerrienet E, et al. Prediction of treatment failure using 2010 World Health Organization Guidelines is associated with high misclassification rates and drug resistance among HIV-infected Cambodian children. Clinical Infectious Diseases. 2012;55(3):432–40.

44. Khienprasit N, Chaiwarith R, Sirisanthana T, Supparatpinyo K. Incidence and risk factors of antiretroviral treatment failure in treatment-naïve HIV-infected patients at Chiang Mai University Hospital, Thailand. AIDS research and therapy. 2011;8(1):42.

45. Soria A, Porten K, Fampou-Toundji J, Galli L, Mougnutou R, Buard V, et al. Resistance profiles after different periods of exposure to a first-line antiretroviral regimen in a Cameroonian cohort of HIV type-1-infected patients. Antiviral therapy. 2009.

46. Okoye AA, Picker LJ. CD 4+ T-cell depletion in HIV infection: mechanisms of immunological failure. Immunological reviews. 2013;254(1):54–64.

47. Endalamaw A, Tezera N, Eshetie S, Ambachew S, Habtewold TD. Adherence to Highly Active Antiretroviral Therapy Among Children in Ethiopia: A Systematic Review and Meta-analysis. AIDS and Behavior. 2018:1–11.

48. Giday A, Shiferaw W. Factors affecting adherence of antiretroviral treatment among AIDS patients in an Ethiopian tertiary university teaching hospital. Ethiopian medical journal. 2010;48(3):187–p94.

49. Markos E, Worku A, Davey G. Adherence to ART in PLWHA and Yirgalem Hospital, South Ethiopia. Ethiopian Journal of Health Development. 2008;22(2):174–9.

